# Jointly Optimized Spatial Histogram UNET Architecture (JOSHUA) for Adipose Tissue Segmentation

**DOI:** 10.1101/2021.11.22.469463

**Authors:** Joshua K. Peeples, Julie F. Jameson, Nisha M. Kotta, Jonathan M. Grasman, Whitney L. Stoppel, Alina Zare

## Abstract

**Objective:** We quantify adipose tissue deposition at surgical sites as a function of biomaterial implantation.

**Impact Statement:** To our knowledge, this study is the first investigation to apply convolutional neural network (CNN) models to identify and segment adipose tissue in histological images from silk fibroin biomaterial implants.

**Introduction:** When designing biomaterials for the treatment of various soft tissue injuries and diseases, one must consider the extent of adipose tissue deposition. In this work, we implant silk fibroin biomaterials in a rodent subcutaneous injury model. Current strategies for quantifying adipose tissue after biomaterial implantation are often tedious and prone to human bias during analysis.

**Methods:** We used CNN models with novel spatial histogram layer(s) that can more accurately identify and segment regions of adipose tissue in hematoxylin and eosin (H&E) and Masson’s Trichrome stained images, allowing for determination of the optimal biomaterial formulation. We compared the method, Jointly Optimized Spatial Histogram UNET Architecture (JOSHUA), to the baseline UNET model and an extension of the baseline model, Attention UNET, as well as to versions of the models with a supplemental “attention”-inspired mechanism (JOSHUA+ and UNET+).

**Results:** The inclusion of histogram layer(s) in our models shows improved performance through qualitative and quantitative evaluation.

**Conclusion:** Our results demonstrate that the proposed methods, JOSHUA and JOSHUA+, are highly beneficial for adipose tissue identification and localization. The new histological dataset and code for our experiments are publicly available.

## 1 Introduction

### Background

The main goals within biomaterial research for soft tissue regeneration are to develop materials that a) actively promote cellular infiltration into the scaffold and b) degrade at similar rates to *de novo* tissue formation. Biomaterials available for surgeons that are applicable for soft tissue injuries include natural and synthetic materials or a combination of both [1, 2]. Silk fibroin, a protein extracted from *Bombyx mori* silkworm cocoons, is a natural biomaterial that has shown promising clinical translation [3]. Silk fibroin has mechanical, structural, and chemical parameters that can be tuned to allow for a wide scope of final biomaterial formulations [4, 5]. As a result, silk fibroin biomaterials have been used for a variety of different applications. However, silk fibroin biomaterials have been fabricated without concern for adipose tissue deposition within and around the biomaterial. In the treatment of skeletal muscle disorders, an implanted biomaterial that results in excessive adipose tissue deposition would be considered undesirable. As skeletal muscle atrophies, increases in adipose tissue occur, leading to deformation, paralysis, and even death. On the other hand, adipose tissue functions as a protective layer for organs and maintains body contours. Meaning applications for biomaterials that promote adipose tissue deposition exist. Quantifying adipose tissue accumulation for different biomaterial formulations is necessary to engineer optimal biomaterials for particular clinical applications. One method to quantify adipose tissue accumulation is to hand label adipocyte area in histological stained images. Manually labeling adipocytes is time-consuming, tedious, and prone to error. Specifically, inconsistency in labeling (*e.g*., different annotators, fatigue) can also introduce bias when identifying regions of interest.

### Deep learning for histological images

To mitigate human bias and increase efficiency, we can train machine learning models to autonomously identify and quantify the adipose tissue in histological sections. Deep learning, a sub-area of machine learning, has been applied to histological images for different tasks including classification, regression, and segmentation [6, 7]. Segmentation tasks include identifying regions of interest pertaining to cells, nuclei, glands, tissue, and tumors [6]. Current state-of-the-art approaches for semantic segmentation are encoder-decoder models [8, 9]. UNET is an encoder-decoder model that has been used for biomedical (and other) domains [9]. One biomedical application of UNET is for histological image segmentation [6, 10, 11]. UNET has copy and crop paths (*i.e*., skip connections) that fuse information from the beginning (*i.e*., encoder) to the end of the network (*i.e*., decoder) to promote the identification and localization of regions of interest [6, 9]. The copy and crop connections also facilitate improved learning (*i.e*., mitigate the effect of vanishing gradient) since these models are typically trained via backpropagation [12]. Despite these advantages, the UNET model can lead to increased computational cost by producing repetitive and unnecessary features [13]. Additionally, high quality datasets are needed to train and evaluate deep learning approaches for histological image segmentation [6].

### Attention mechanisms

In machine learning, attention mechanisms have been introduced to improve the performance of models for different applications including natural language processing and semantic segmentation [8, 14, 15]. The motivation behind these approaches is to better train the model to focus on the most relevant and important features in the data [8, 14, 16]. Attention mechanisms have been integrated into UNET to improve segmentation performance (*e.g*., Attention UNET) [13]. Generally, attention mechanisms learn how to weight (*i.e*., place importance) the input features of the data to encode information such as positions in a sequence [16]. However, attention mechanisms can increase computational costs by increasing the number of learnable parameters in the model. We hypothesize that a simpler (*i.e*., fewer parameters) attention-inspired fusion approach will lead to improved performance while reducing computational constraints (more details in Section 4.3).

### Problem statement

Determining adipose tissue accumulation around and within degrading biomaterial-based implants allows engineers and scientists to create biomaterials for specific clinical applications. Previous methods to identify adipose tissue in histological images are tedious, laborious, and susceptible to human bias and error. UNET is a rapid, efficient, and high-throughput method that can be used to quantify adipose tissue. As currently constructed, UNET does not directly encode statistical texture information which has improved performance for semantic segmentation [17–19]. To improve the representation of information throughout the convolutional neural network (CNN), statistical texture features can be used to better characterize the spatial distributions of the data, potentially leading to improved performance as shown in other works that incorporate statistical texture information [7, 17, 18, 20].

In this work, acellular silk fibroin lyophilized sponges were implanted into the lateral pockets of Sprague Dawley rats. After 1-, 2-, 4-, and 8-weeks post-surgery, the silk sponges and overlaying tissue were excised. Samples were prepped and stained with hematoxylin and eosin (H&E) and Masson’s Trichrome. To improve the identification and quantification of adipose tissue, we proposed to extract statistical texture features through the use of histogram layer(s) [21] integrated into the baseline architecture, termed Jointly Optimized Spatial Histogram UNET Architecture (JOSHUA). Additionally, we hypothesized that statistical texture features could also be used as an attention approach to weight important information in the network in order to improve the context learned by each model during training, termed JOSHUA+. The overall framework of this study is shown in Figure 1. Our contributions to the field include:

- Novel incorporation of histogram layers to extract statistical texture and spatial information to improve segmentation performance
- New dataset for the community to evaluate histological images for adipose tissue segmentation
- Simple yet effective attention-inspired approach to reduce parameters of the model while achieving comparable and/or better performance
- JOSHUA+ accurately identifies differences in adipose tissue accumulation in biomaterial formulations over time
- JOSHUA+ decreases the time to find adipose tissue area by 200% compared to manually annotating

**Figure 1:**
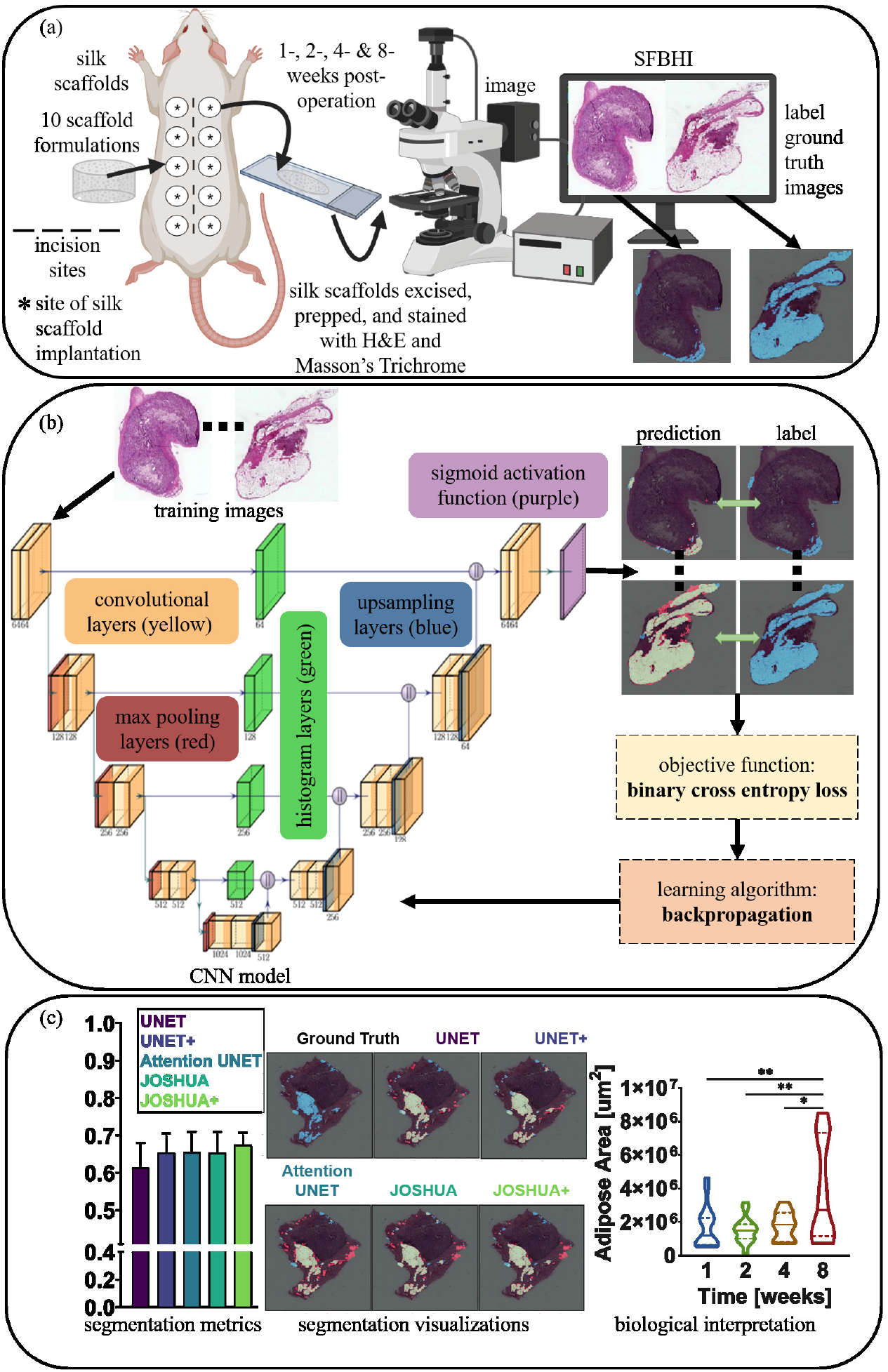
Our overall process has three components: data collection, processing, and analysis as shown in Figures 1a through 1c, respectively. (a) For data collection, acellular silk fibroin lyophilized sponges of varying formulations were subcutaneously inserted into the lateral pockets of Sprague Dawley rats. After 1-, 2-, 4-, and 8-weeks post-surgery, the silk sponges and overlaying tissue were excised. Samples were prepped and stained with H&E and Masson’s Trichrome. (b) The next step (*i.e*., data processing) trains our proposed model through *k*-fold cross validation. (c) Once training is completed, we can use our machine learning models to quickly segment and quantify the adipose tissue for new samples. We then perform data analysis to connect the model outputs to meaningful biological interpretations.

## 2 Results and Discussion

### 2.1 Data Selection using UNET

For machine learning approaches, it is necessary to understand how splitting the data into training and validation sets influence the results of the model using standard segmentation assessment metrics as shown in Figure 2 and Supplemental Table S1. To do this, we evaluated the impacts of time and biomaterial conditions to determine how the data influenced the performance of the model. Ideally, we need a model that can be applied to analyze data from any time point and any silk biomaterial formulation. We investigated five training and validation splits of our novel *Silk Fibroin Biomaterial Histology Images* (SFBHI) dataset with the baseline UNET model: Random, Stratified 5-fold (Time), Stratified 5-fold (Condition), 4-fold with Time, and Validate on week 8. For our SFBHI dataset, we developed *ground truth* for each image by labeling each pixel as either *adipose tissue* or *background*. The ground truth is then used to train the machine learning model to produce predictions based on the input images. We used six metrics to assess segmentation performance on the validation images: Sørensen–Dice coefficient, referred to as dice coefficient, overall intersection over union (IOU), adipose tissue IOU, precision, recall, and specificity. Each metric ranges between 0 and 1. A value of 0 means there is no overlap between ground truth and model prediction while a value of 1 means the ground truth and model prediction overlap perfectly. Details relating to the computation of the segmentation metrics are in Section 4.1.

**Figure 2:**
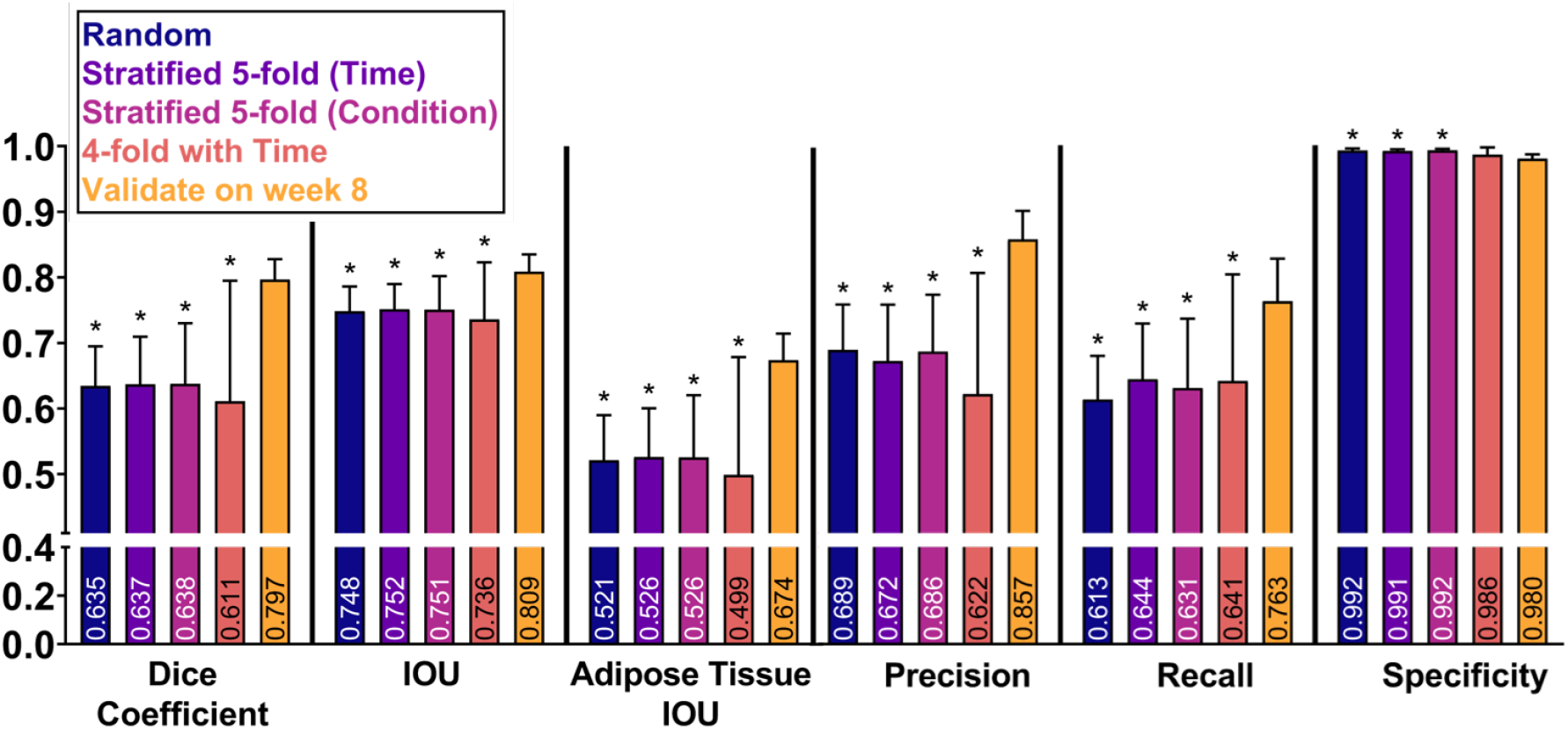
Dice coefficient, IOU, adipose tissue IOU, precision, recall, and specificity metrics for data splits, random, stratified 5-fold (time), stratified 5-fold (condition), 4-fold with time, and validate on week 8, on validation images in SFBHI dataset trained on UNET with weighted binary cross entropy. Data are shown as mean ± SD. One-way analysis of variance, followed by Dunnett’s multiple-comparison test was computed. Asterisks [*] indicate significant differences as compared to the *validate on week 8* data split (p < 0.05).

The five training and validation splits of our novel SFBHI dataset impacted the performance of the baseline UNET model (Figure 2). As detailed in Section 4.1, we used k-fold cross validation. Supplemental Tables S2–S7 detail the percentages of each condition and week used for every data split (Random, Stratified 5-fold (Time), Stratified 5-fold (Condition), 4-fold with Time, Validate on week 8). We found that validating the UNET model on week 8 SFBHI images showed a statistically significant improvement in performance for all metrics except for specificity (Figure 2). Week 8 has the most accumulation of adipose tissue, and as a result, these images are easier for the model to segment and improve metrics such as intersection-over-union (IOU). These models may possibly fail to generalize and perform well when presented with more difficult images (*e.g*., sparse areas of fat accumulation). This point is further validated by the drop in performance when performing 4-fold cross validation with Time. For three of the four folds, week 8 images are used as training data. However, the overall average is significantly less than validating on week 8 only. Since the model used the “easier” training samples, UNET did not perform as well on the more difficult images with small regions of adipose tissue.

Another interesting observation is that the results of random and stratified (*i.e*., time, condition) 5-fold cross validation are comparable to one another (Figure 2). The Stratified 5-fold cross validation approach considered the biologically-meaningful labels while the Random 5-fold cross validation did not. The dataset is relatively balanced across the number of images for each time or condition. In the case of possible data imbalance for future applications, it may be important to leverage biological information (*e.g*., time and condition) to properly train and validate the model. When comparing the results of stratified 5-fold with time and condition, the model is robust to the use of either biological label. As a result, the model can learn a meaningful mapping between the input and output to identify adipose tissue in either case of labeling.

### 2.2 Model Comparisons

In practice, we want to deploy the most robust model that will generalize well to both dense and sparse accumulations of adipose tissue. Therefore, we selected the random 5-fold cross validation split to compare the baseline UNET model to our proposed JOSHUA, attention-inspired variants (JOSHUA+ and UNET+), and Attention UNET [13] models. The results of the model comparisons are shown in Figure 3 and Table S8. Example qualitative results of segmenting the validation images using our trained models are shown in Figure 3a. Silk-collagen I (s-c), silk+VEGF_*S*_ (s+VEGF_*S*_), and silk biomaterials were selected as “easy” data samples with robust adipose presence whereas additional training was also done on adipose-poor samples such as silk-heparin+VEGF_*S*_ (s-h+VEGF_*S*_) and s-c-h-VEGF. All models performed well on histological images with large areas of adipose tissue. We observed that the histogram-based models are able to better capture small areas of accumulated fat tissue in comparison to the UNET model as shown on the image from week 8 and condition s-c (Figure 3a, third row). JOSHUA and JOSHUA+ also reduce the number of false positives in comparison to Attention UNET for the image from week 8 and condition s-h+VEGF_*S*_ (Figure 3a, first row). The histogram layer(s) integrated into the models demonstrate the effectiveness of statistical texture features for biomedical applications as shown throughout the literature [7, 18, 20].

**Figure 3:**
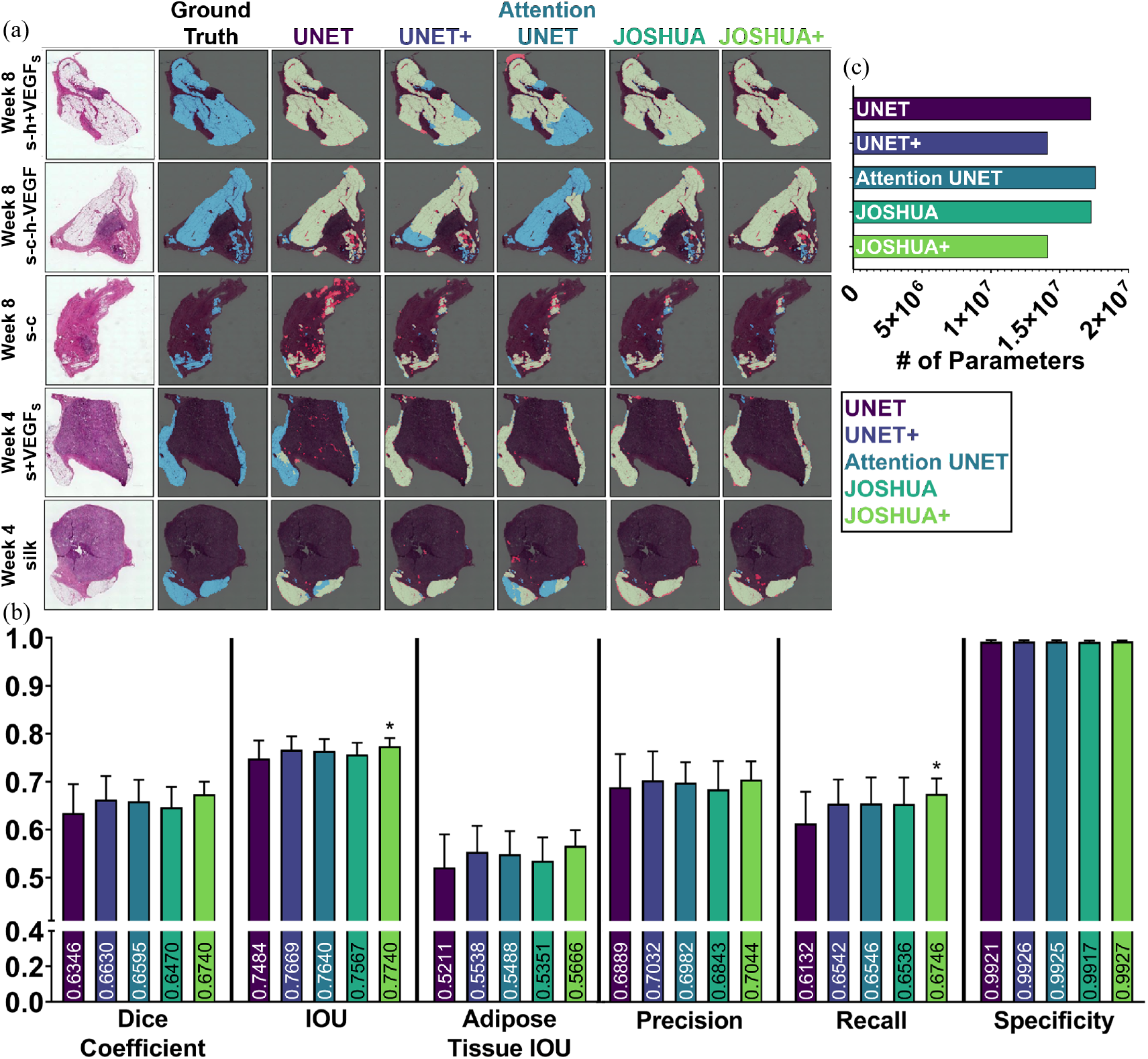
(a) Example segmentation results from each model on the SFBHI dataset. The first column displays the input image and the corresponding ground truth label for the input image is shown in the second column. The remaining columns display the output from each model in comparison to the ground truth. Blue pixels correspond to the ground truth and red pixels correspond to the predicted output. Light green pixels indicate that both the ground truth and predicted output agree. (b) Dice coefficient, IOU, adipose tissue IOU, precision, recall, and specificity metrics for each model for SFBHI dataset. As shown here, the histogram models (JOSHUA/JOSHUA+) and UNET+ improve segmentation results compared to the baseline UNET and Attention UNET [13] models. Metrics are shown as mean ±SD. One-way analysis of variance, followed by Dunnett’s multiple-comparison test were computed. Asterisks [*] indicate significant differences as compared to UNET (p < 0.05). (c) We also evaluated the number of learnable parameters for each CNN model. The attention-inspired variants have approximately 18% fewer parameters than their counterparts.

When observing the mean of each metric, JOSHUA consistently achieves a higher value except for precision and specificity. This indicates that the addition of histogram layer(s) into a network provides more informative features to improve segmentation performance. The metric with the largest difference between JOSHUA and UNET is recall. Recall informs how well each model can identify adipose tissue in our images, as detailed in Section 4.1. This observed difference in recall further verifies our hypothesis that statistical texture features are beneficial for capturing adipose tissue accumulation. The average precision and specificity of JOSHUA are slightly lower than the baseline UNET model, indicating a possible overestimation of adipose tissue due to increased sensitivity (*i.e*., recall) with the addition of histogram layers. JOSHUA could be further improved in later work by increasing the local window size of the histogram layers to improve the discrimination of adipose tissue and background. By increasing the window size, the model can capture more pixel and/or feature values to estimate distinct distributions between the background and adipose tissue.

We found improved performance with the attention-inspired variants of each model, UNET+ and JOSHUA+. Our new method of fusing features from the encoder and decoder components of the architecture encourages joint agreement between features at multiple levels in the network. By encouraging this joint agreement, we improved the mean value for each metric in comparison to the baseline UNET model. This result demonstrates that our fusion approach is beneficial for both the positive (*i.e*., adipose tissue) and negative (*i.e*., background) classes. JOSHUA+ is the only model to achieve a statistically significant difference for two metrics, IOU and recall. For overall IOU, JOSHUA+ improves the identification and localization of not only the adipose tissue but also the background information. JOSHUA+ also shows that statistical texture features are more informative as an attention mechanism in comparison to the convolutional feature maps of UNET+. UNET+ and JOSHUA+ have a higher average than Attention UNET for most metrics. Our proposed attention-inspired models have fewer learnable parameters than Attention UNET as shown in Figure 3c. As a result, the storage requirements for the trained attention models will be reduced and the fusion operation (*i.e*., elementwise multiplication) is quicker to compute than the concatenation approach of UNET, Attention UNET, and JOSHUA.

Example qualitative results of segmenting the validation images using our trained models are shown in Figure 3a. All models perform well on large areas of adipose tissue in the histological images. We observe that the histogram-based models are able to better capture small areas of accumulated fat tissue in comparison to the UNET model as shown in the image from week 8 and condition s-c (Figure 3a, third row). JOSHUA and JOSHUA+ also reduce the number of false positives in comparison to Attention UNET for the image from week 8 and condition s-h+VEGF_*s*_ (Figure 3a, first row). The visualizations are shown to demonstrate the ability of statistical texture features to refine the identification and segmentation of adipose tissue. To provide more insight into the incorporation of statistical textures, we computed the color histograms of the SFBHI dataset in Supplemental Figure S2. The distributions of the adipose tissue and background pixels are very different from one another (*i.e*., little overlap). As a result, all of the models will be able to correctly classify most pixels in the images which led to limited statistical differences for most metrics in Figure 3b. However, there is some overlap with some of the pixel intensity values. In these instances, the statistical texture features are needed to improve performance on these more difficult samples in the data as shown in the segmentation results in Figure 3a.

### 2.3 Application of Models to Benchmark Histological Dataset

To demonstrate the effectiveness of the proposed model to be generalized to related segmentation tasks, we evaluated each model with a benchmark dataset, Gland Segmentation in Colon Histology Images (GlaS), for cancerous histological segmentation. The results of the experiments are shown in Figure 4 and Table S3.

**Figure 4:**
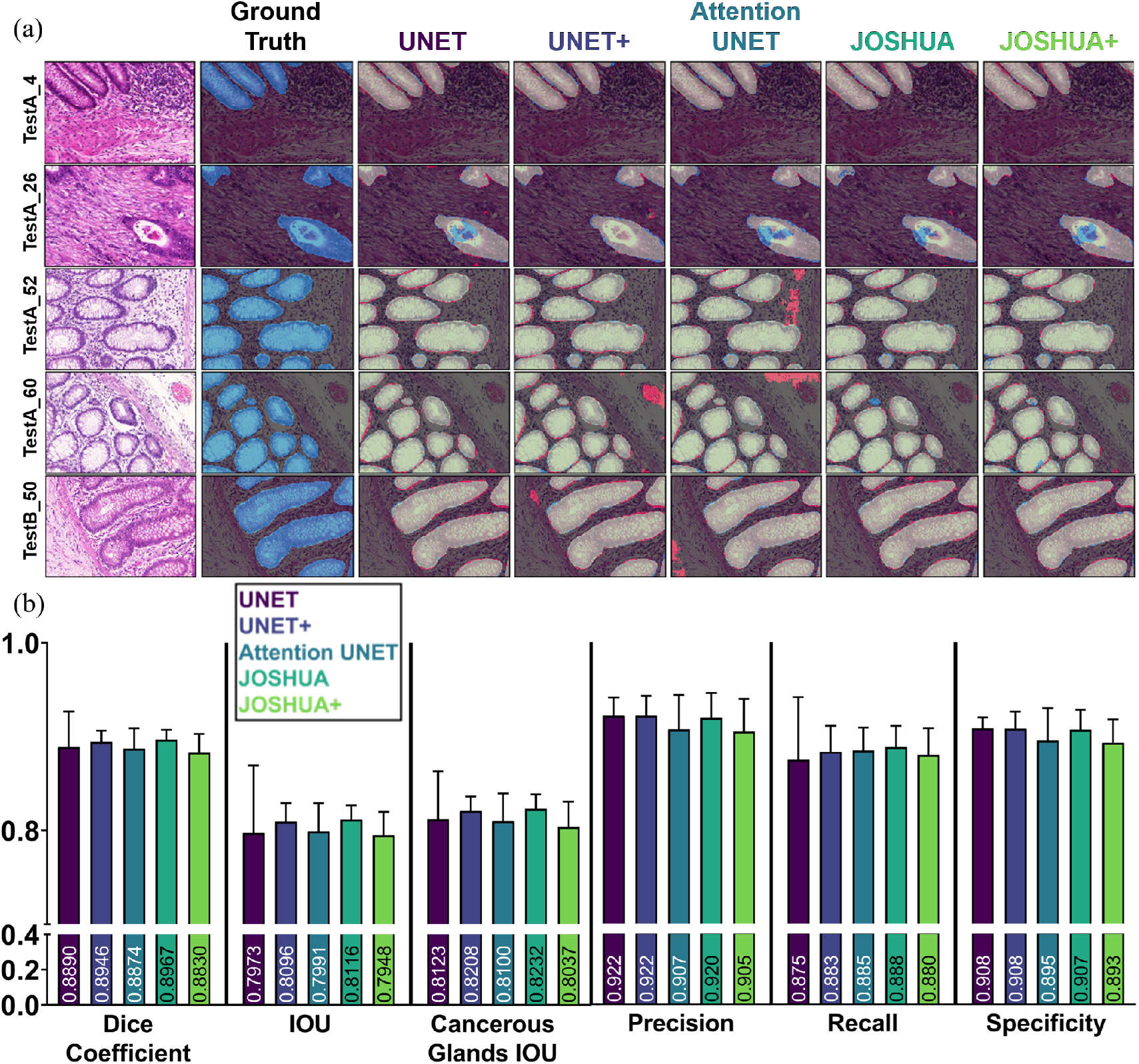
(a) Example segmentation results from each model on the GlaS dataset. The first column displays the input image and the ground truth label for the input image is shown in the second column. The remaining columns display the output from each model in comparison to the ground truth. Blue pixels correspond to the ground truth and red pixels correspond to the predicted output. Light green pixels indicate that both the ground truth and predicted output agree. As shown here, all models perform comparatively well for this dataset. In some instances, the models with the histogram layers (*i.e*., JOSHUA and JOSHUA+) reduce the false positives identified by the other models. (b) Dice coefficient, IOU, adipose tissue IOU, precision, recall, and specificity metrics for each model for GlaS dataset. Metrics are shown as mean ±SD. One-way analysis of variance, followed by Dunnett’s multiple-comparison test were computed. Asterisks [*] indicate significant differences as compared to UNET (p < 0.05).

All models perform comparatively with the baseline UNET architecture with no significant differences (Figure 4b). JOSHUA achieves the highest average for most metrics except for specificity. JOSHUA, JOSHUA+, and UNET+ improved the sensitivity (*i.e*., recall) over the UNET model. There is a large improvement in each metric (except specificity) for every model in comparison to the SFBHI dataset. The proposed approaches (JOSHUA/JOSHUA+) also have less variability (*i.e*., small standard deviations) than the UNET model across the metrics.

Overall, JOSHUA achieves the highest average for most metrics except for specificity. The GlaS dataset has a “zoomed in” view of the histological images resulting in reduced background from the histological slides. Therefore, we see a large improvement in each metric (except specificity) for every model in comparison to the SFBHI dataset due a reduction in false positives as a result of the slide background. An important note here is that JOSHUA, JOSHUA+, and UNET+ improved the sensitivity (*i.e*., recall) over the UNET model. This result further validates that our histogram layer(s) and fusion approaches better identify the class of interest (*i.e*., cancerous or adipose tissue). Our proposed approaches also have less variability (*i.e*., small standard deviations) than the UNET model across the metrics. We also calculated the color histograms of the GlaS dataset in Supplemental Figure S3. The distributions of the cancerous tissue and background pixels are more similar in comparison to the distributions of adipose tissue and background pixels of the SFBHI dataset. For the GlaS dataset, the concatenation approach of JOSHUA was more effective than the elementwise multiplication of JOSHUA+ in terms of achieving better segmentation metrics. This result indicated that if the class of interest (*i.e*., adipose or cancerous tissues) has a more similar distribution of values to the background, JOSHUA will be more effective than JOSHUA+.

Qualitatively, we see similar results to the SFBHI dataset. Each model performs well for large cancerous regions of the image. However, the UNET+ and Attention UNET model detect more falsely identified cancerous glands for images in the third and fourth row of Figure 4a). For the second row, all models (except for UNET+) have difficulty with detecting the central area of the cancerous gland on the bottom right hand side of the image. In comparison to the other images, this gland has the most abrupt changes in terms of statistical texture and color information. As stated in the SFBHI section, the window size of the histogram layer(s) can be modified to better characterize the distribution of cancerous and non-cancerous glands to account for images with similar characteristics. The UNET+ model has the fewest number of parameters. This can lead to improved generalization ability in distinct cases such as the “TestA_26” image.

### 2.4 Applying Model on New Images

To investigate the use of JOSHUA+ to identify the differences in adipose tissue accumulation based on varying biomaterial formulations over time, we took the best JOSHUA+ model (*i.e*., random initialization three and fold three) to plot the adipose area estimation while considering condition and time (Figure 5). In addition to the original 117 images, we applied our model to 465 new images that were not labelled. Figure 5a shows segmentation results from both H&E and Masson’s Trichrome stained holdout images. We observed that JOSHUA+ falsely identified adipose tissue for some images (*i.e*., week 2 H&E, week 1 Masson’s Trichrome). We expected this would occur because lyophilized silk fibroin sponges can have spherical pores that can be mistaken for spherical adipocytes. JOSHUA+ could be used as a “pre-processing” step to initially identify adipose tissue (or other regions of interest) in tandem with human annotators to ease the burden of labeling images. Figure 5b is a graph of adipose area over time for the representative images shown in Figure 5a. The area of adipose tissue at week 8 was significantly higher than the adipose area at weeks 1, 2, or 4 adipose the for s-c+VEGF_*S*_ biomaterial formulation. Figure 5c shows the difference in adipose area across 10 biomaterial formulations for week 4. The s-c condition is the only biomaterial formulation to show a significant increase in adipose area at week 4 compared to silk biomaterials. These are important findings as we can produce statistical differences in adipose area over time with varying biomaterial formulations. Future work is needed to determine adipose tissue area with respect to section and scaffold areas to identify biological interpretations of adipose tissue as a function of time and biomaterial formulation.

**Figure 5:**
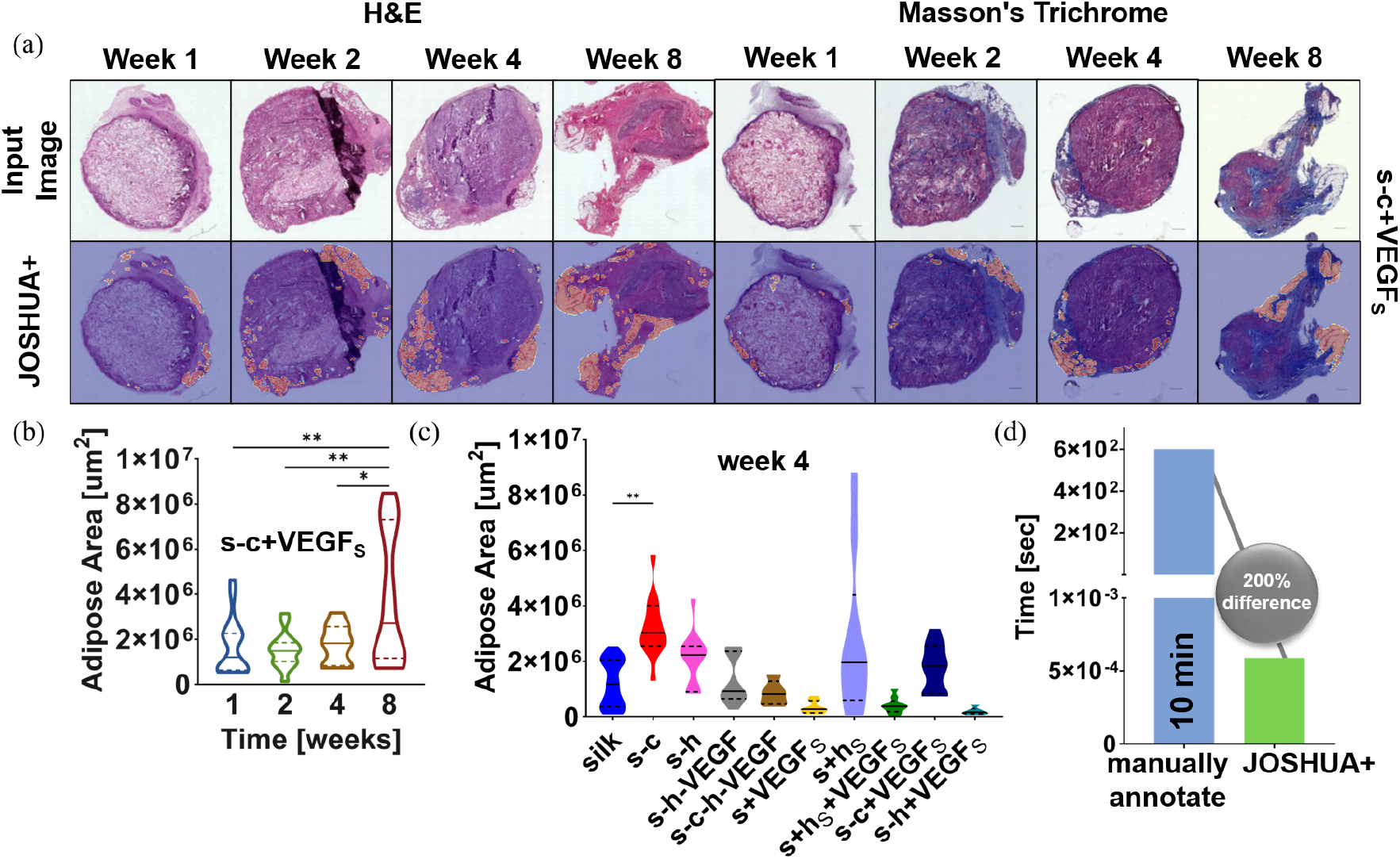
(a) SFHBI holdout images and their segmentation results using JOSHUA+. H&E images and their corresponding segmentation results are shown in columns 1-4. Masson’s Trichrome images and their corresponding segmentation results are shown in columns 5-8. (b) Quantification of adipose area over time for s-c+VEGF_*S*_ biomaterial formulation. (c) Adipose area across 10 biomaterial formulations for week 4. (d) The time to manually annotate the adipose area or use JOSHUA+ for one image. There is a 200% difference in time to use JOSHUA+ rather than manually labeling and quantifying adipose area. For Figure 5b & 5c, data are shown with a solid line representing the median and the two dashed lines representing the 25^*th*^ and 75^*th*^ percentile quartiles. Kruskal-Wallis test, followed by Dunn’s multiple-comparison test was computed. Asterisks [*] indicate significant differences (p < 0.05).

Figure 5d shows the time in seconds to quantify the adipose area for one image for both manually labeling and using JOSHUA+. This new strategy enables the processing of much larger datasets compared to the existing methods with significantly faster computation speed by leveraging recent advances in deep learning. A limitation of our current work is that the dataset is relatively small for machine learning approaches. As a result, additional images would be beneficial to train and verify the model for improved performance. However, this data set is relatively large for biomaterial scientists and engineers. Another constraint is that the model was trained using ground truth labels from one annotator. These labels will have some bias; therefore, our method will also benefit by using labels from multiple expert annotators to further improve the robustness and accuracy of the models. In summary, we successfully trained and validated the model with a subset of the data (117 images) and then used JOSHUA+ to analyze a full data set of 465 images.

## 3 Conclusion

In this work, we quantified adipose tissue deposition at surgical sites as a function of biomaterial composition and the time post-implantation. Acellular silk fibroin lyophilized sponges were implanted into the lateral pockets of Sprague Dawley rats. After 1-, 2-, 4-, and 8-weeks post-surgery, the silk sponges and overlaying tissue were excised. Samples were prepped and stained with H&E and Masson’s Trichrome. We used CNN models with novel spatial histogram layer(s) to identify and segment regions of adipose tissue in the histological images, allowing for the determination of the optimal biomaterial formulation. We compared our proposed method, JOSHUA, to the baseline UNET model and an extension of the baseline model, Attention UNET, as well as to versions of the models with a supplemental attention-inspired mechanism (JOSHUA+ and UNET+). Our results demonstrate that the proposed methods, JOSHUA and JOSHUA+, are highly beneficial for adipose tissue identification and localization within and around silk fibroin lyophilized sponges. To our knowledge, this study is the first investigation to apply CNN models to identify and segment adipose tissue in histological images from silk fibroin biomaterial implants. It is also the first step in developing automated analysis to guide the formulation of smart biomaterials that can meet the needs of clinicians, surgeons, and patients. Future work aims to determine the components of the biomaterial formulation that drive adipose tissue deposition post-surgical implantation. Additionally, our machine learning process could be further improved by tuning hyperparameters (*e.g*., window size and the number of bins) and exploring weakly supervised learning approaches to reduce the cost of acquiring precise, pixel-level ground truth labels.

## 4 Materials and Methods

### 4.1 Experimental and Technical Design

#### Description of Datasets

In the SFBHI and GlaS datasets, there are 117 and 165 images respectively. The SFBHI dataset has images at various scales and illuminations as shown in Figure S1a. Ground truth labels were created for our SFBHI dataset using Hasty [22], an image annotation software. Regions of adipose tissue accumulation in 117 histological images of various silk scaffold conditions were labeled by hand using the software’s semantic segmentation feature. An extension of our SFBHI dataset had a total of 465 images that were unlabeled and had two stainings: H&E and Masson’s Trichrome. These new images were used to evaluate the trained model (*i.e*., JOSHUA+) for quantifying and analyzing adipose tissue across time and biomaterial formulation.

#### Evaluation and Training Protocol

For both datasets, we adopt a five-fold cross validation strategy. There were a total of 94 (93 for folds three through five) training images and 23 (94 for folds three through five) validation images for SFBHI in each fold. The training (67 images for folds one through four, 72 for fold five) and validation (18 images for folds one through four, 13 for fold five) data splits from Rony et. al. [23] were used for the GlaS dataset. For each dataset, we ran three runs of random initialization for a total of 15 experimental runs. The results shown for SFBHI are averaged across the validation folds for each model while the GlaS results are average performance metrics on the holdout test evaluated by each model trained and validated on each fold. Convolutional models are dependent on scale [24], so we used bilinear interpolation to resize the SFBHI images to be the same resolution (256 × 256). The GlaS images did not need to be resized since all images were the same resolution (775 × 522). We compare a total of five models: baseline UNET [9], UNET+, Attention UNET [13], JOSHUA, and JOSHUA+.

To improve the performance and robustness of each model, we followed a similar data augmentation procedure to Rony et. al. [23]. The images are randomly flipped horizontally (p =0.5) and rotated by 90°increments (*i.e*., 0°, 180°, and 270°). Random color jittering (*i.e*., changes in brightness, contrast, and saturation at *p* = 0.5 and hue at *p* = 0.05) is also incorporated in the training images. The images are also iterated over eight times within each mini-batch to further artificially increase the size of the training dataset. Random crops of 416 × 416 are extracted during training for the Glas dataset. The data splits and random initialization (three random seeds) of each model were fixed so we could do a fair comparison across the different architectures. Each model was trained for 150 epochs while using Adam [25] optimization with an initial learning rate of 0.001, weight decay 1 × 10^-8^, and the gradient values were clipped at 0.1. Early stopping was also implemented to prevent overfitting on the training data. The early stopping epochs were set to 10 for SFBHI and 20 for GlaS. Batch sizes of four and eight were used for GlaS and SFBHI respectively. Experiments were run on two NVIDIA GeForce RTX 2080TI GPUs. We used binary cross entropy as the loss function to train the models. The SFBHI dataset is heavily imbalanced as some images do not contain much adipose tissue. To counter this, we used weighted binary cross entropy and set the weight in the loss function to three (determined empirically).

In our work, we looked at using stratified and non-stratified cross validation (Section 2.1). For stratification, the division of the data is based on maintaining a similar distribution of labels in both training and validation folds. For our histological images, we have two associated labels: time (1-, 2-, 4-, and 8-weeks post-surgery) and condition (silk, silk-collagen I, silk-heparin, silk-heparin-VEGF, silk-collagen I-heparin-VEGF, silk+VEGF_*S*_, silk+heparin_*S*_, silk+heparin_*S*_+VEGF_*S*_, silk-collagen I+VEGF_*S*_, silk-heparin+VEGF_*S*_). We first used a data split based on a random partition of 5-fold cross validation that did not consider the global labels (*i.e*., time and condition) associated with each image as in the stratified examples. The distribution of global labels are shown in Supplemental Tables S2 and S3. Using stratified cross validation, we had two data splits: stratified 5-fold (time) and stratified 5-fold (condition) (see Supplemental Tables S4 and S5, respectively). The final two data splits used the time information to divide the data based on the weeks after surgery. In this instance, each fold represents a specific week (*e.g*., train on 1-, 2-, and 4-weeks, validate on week 8). Since there are only four distinct time points, 4-fold cross validation was used for this data split (*i.e*., 4-fold with time) (Supplemental Table S6). Lastly, we also experimented with only validating on week 8 images (Supplemental Table S7). By using this data split, we wanted to evaluate if the model(s) could learn sequentially (*i.e*., train on earlier weeks, predict later weeks).

#### Evaluation Metrics

There are several common metrics to assess semantic segmentation performance. The first is pixel accuracy, which measures how well the model labels each pixel to the corresponding class. For our segmentation work, we have a two possible classes: positive (*i.e*., adipose and cancerous tissue) and negative (*i.e*., background). With these two classes, we have four possible outcomes: true positive (*i.e*., positive samples correctly identified as positive), false positive (*i.e*., negative samples incorrectly identified as positive), true negatives (*i.e*., negative samples correctly identified as negative), and false negatives (*i.e*., positive samples incorrectly identified as negative). Pixel accuracy is defined as the sum of true positives (TP) and negatives (TN) divided by the sum of all pixels in the image as shown in Equation 1:

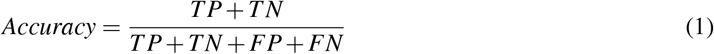

where FP is false positive and FN is false negative. However, pixel accuracy is not an optimal metric to use when there is a severe class imbalance (such as with the SFBHI dataset for adipose tissue vs. background). Therefore, we need other metrics to provide more insight into each model’s performance. Three other measures are precision, recall, and specificity. Precision quantifies the correctly identified positive samples, recall (*i.e*., sensitivity) is how well the model can identify positive samples, and specificity is how well the model identifies true negative samples. Precision, recall, and specificity are defined in Equations 2, 3, and 4 respectively:

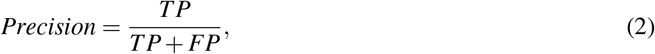

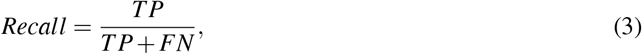

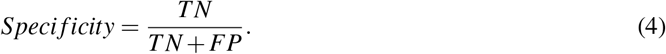

There is a trade off between precision and recall (*i.e*., as one increases, the other will decrease). To find the balance between the two measures, the F1 measure can be used. For binary problems, the F1 score is equivalent to the Dice coefficient [26] and is shown in Equation 5:

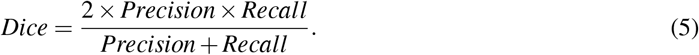

A related measure to the Dice score is Intersection Over Union (IOU), also known as the Jaccard index. IOU is defined as the intersection of the model prediction, *Ŷ*, and the true label, *Y*, divided by the union of the two sets:

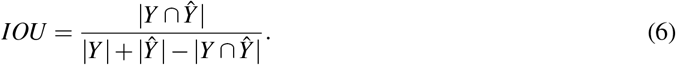

For IOU, we report the results on the positive class or regions of interest as well as the overall IOU (background and positive classes).

#### Quantification of Adipose Tissue

In addition to evaluating segmentation performance, we also wanted to compute the measurements of adipose tissue found by the machine learning models. To train and evaluate the models, we needed to resize the images to a smaller size as stated previously. In order to scale the amount of adipose tissue area (*μm*^2^) in the reduced images, we used the following computation in Equation 7 where the FA_*full*_ is the full resolution fat pixel area, FA_*down*_ is the downsampled fat pixel area, IA_*full*_ is the full resolution image pixel area, IA_*down*_ is the downsampled image pixel area, and RL is the reference length from the Keyence^®^ Analysis software 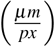:

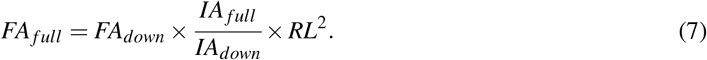

### 4.2 Dataset Collection: Silk Fibroin Biomaterial Histology Images (SFBHI)

#### Silk Fibroin Extraction

As previously described, silk fibroin solution was prepared through isolation from *Bombyx mori* silkworm cocoons [5]. Five grams of silk cocoons were degummed in 2 L of boiling 0.02 M sodium carbonate (Catalog No. 451614, Sigma Aldrich, St. Louis, MO) for 30 minutes in order to isolate pure silk fibroin protein. After degumming, the silk fibroin protein was left to air dry in the fume hood for at least 48 hours. The dried silk fibroin protein was then solubilized by denaturation in 9.3 M aqueous lithium bromide (Catalog No. 213225, Sigma Aldrich, St. Louis, MO) at 60°C for 4 hours. Using a 3.5 kDa molecular weight cutoff dialysis membrane tubing (Catalog No. 08-670-5C, Spectrum™ Spectra/Por™ 3 RC Dialysis Membrane Tubing, 3,500 Dalton MWCO, Thermo Fisher Scientific, Waltham, MA), the solution was dialyzed in deionized water to remove the lithium bromide ions. This produces a solubilized silk solution that was centrifuged twice (>2000 g, 20 min, 4°C) to remove insoluble particles. The silk solution concentration was determined by placing a wet volume of the silk solution in the oven at 60°C and massing the final dry weight. This protocol resulted in a final solution between 5–7% weight per volume (wt./v, 0.05 – 0.07 g/mL) silk solution. Silk solutions were stored at 4°C for up to 3 weeks prior to use in making silk sponges.

#### Type I Collagen Isolation

All animal experiments were executed under protocols approved by both Tufts University’s and University of Florida’s Institutional Animal Care and Use Committees and in accordance with the Guide for the Care and Use of Laboratory Animals (NIH, Bethesda MD). As previously described [27], ready-to-use reconstituted type I collagen was prepared. Briefly, rat tail type I collagen was isolated from the tails of adult Sprague Dawley rats. Following lyophilization, the resulting collagen sponge-like material was solubilized in 0.02 N acetic acid.

#### Scaffold Formation

Two methods of silk scaffold formation were pursued: pre-fabrication method and post-fabrication method. The pre-fabrication method starts with incorporating heparin (Catalog No. H3149-250KU), heparin sodium salt from porcine intestinal mucosa (250KU, Sigma Aldrich, St. Louis, MO, 100 μg/mL), type I collagen (200 μg/mL), and vascular endothelial growth factor (VEGF) (Catalog No. 100-20, Recombinant Human VEGF165, Peprotech, Rocky Hill, NJ, 0.1 μg/mL). A mixture of heparin, collagen I, and VEGF was created prior to being added to diluted aqueous silk solution (3% weight per volume (wt./v, 0.03 g/mL). Following published protocols [4], the silk solution was poured into wells of a 6 well plate and frozen in −80°C freezer overnight, forming isotropic silk scaffolds. Sponges were lyophilized at −80°C and 0.185 mbar for 5-7 days (FreeZone 12 Liter −84°C Console Freeze Dryer, Labconco, Kansas City, MO) post-freezing. To make the sponge-like scaffolds water insoluble, the scaffolds were water annealed for 12 hours at room temperature to induce *β*-sheet formation (silk fibroin protein crystallization) [28, 29]. Sponge-like scaffolds were cut using a 6 mm diameter biopsy punch to a create cylinder 3 mm in height. The post-fabrication method is described by pouring the diluted aqueous silk alone into wells of a 6 well plate and freezing the solution in −80°C freezer overnight. This is again followed by lyophilizing the silk sponge at −80°C and 0.185 mbar for 5-7 days (FreeZone 12 Liter −84°C Console Freeze Dryer, Labconco, Kansas City, MO). These silk sponges were also made water insoluble by water annealing for 12 hours at room temperature to induce *β*-sheet formation (silk fibroin protein crystallization) [28, 29]. Prior to trimming the scaffolds, VEGF and heparin were solubilized where VEGF_*S*_ and heparin_*S*_ designate soluble VEGF and heparin added post-fabrication, respectively. Scaffolds were then soaked in solubilized solutions of VEGF_*S*_ and/or heparin_*S*_ for 30 minutes. Sponge-like scaffolds were cut using a 6mm diameter biopsy punch to a create cylinder 3 mm in height.

#### *In vivo* Subcutaneous Procedures

All procedures were conducted under animal protocols approved by Tufts Institutional Animal Care and Use Committee. All animals used in this study were 7-8-week-old Sprague Dawley rats (~225 g, Charles River Laboratories, Wilmington, MA). Following general anesthesia of oxygen and isoflurane, the back of the rat was shaved, and the skin was prepped using three sets of betadine disinfectant followed by an alcohol wipe. Sterilized acellular silk scaffolds (6 mm diameter x 3 mm height) were subcutaneously implanted in lateral pockets of each rat through up to three small longitudinal incisions made through the skin. The incisions were then closed using surgical clips. Animals were euthanized at 1-, 2-, 4-, and 8-weeks post-surgery. The silk scaffolds (along with the overlying tissue) were then excised and collected for histological examination.

#### Histological Staining of *in vivo* Samples

After a series of graded ethanol and xylene incubations, samples were fixed with 10% phosphate buffered formalin (Thermo Fisher Scientific, Waltham, MA) and embedded in paraffin. Samples were sectioned to 7 – 15 μm thickness and deparaffinized before staining with hematoxylin (Catalog No. SDGHS280, Thermo Fisher Scientific, Waltham, MA) and eosin (Catalog No. SDHT1101128, Thermo Fisher Scientific, Waltham, MA) or Masson’s Trichrome (Catalog No. HT15, Sigma-Aldrich, St. Louis, MO). Hematoxylin stains cell nuclei in a dark purple and blue color while eosin stains the extracellular matrix and cytoplasm pink. Masson’s Trichrome is a multi-color staining technique. Keratin and muscle fibers are stained in red. Collagen and bone are stained in blue or green. Cytoplasm is stained in light red and pink. Cell nuclei are stained in dark brown or black. Samples were embedded in DPX Mountant (Sigma-Aldrich, St. Louis, MO) following staining and dehydration, and then imaged using a Keyence^®^ BZ-X800 series microscope at 20X magnification.

### 4.3 Jointly Optimized Spatial Histogram UNET Architecture (JOSHUA)

The proposed model, Jointly Optimized Spatial Histogram UNET Architecture (JOSHUA), is shown in Figure 6. The model integrates histogram layers [21] in the architecture along the copy and crop connections. By adding the histogram layers in these locations, we are able to include statistical texture information from the encoder and concatenate these features in the decoder to improve the context provided by the “binned” convolutional feature maps. Given *D* number of convolutional features maps of height *M* and width *N* from the encoder, 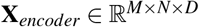, the *K* number of *B*-bin histogram representations of height *R* and width *C*, 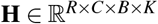, is computed using Equation 8 with a sliding window of size *S* × *T*:

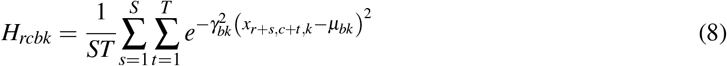

where *μ_bk_* and *γ_bk_* are the bin center and widths, respectively, of the *b^th^* histogram bin for the *k^th^* input feature map. After the convolutional features are binned (mapped to the range of [0, 1]), we aggregate the average counts for each bin across the spatial dimensions of the feature maps to compute the local histograms (*S* = 2, *T* =2). The bin centers and widths of the histogram layers are updated via backpropagation during training in order to improve the representation of the statistical texture features.

**Figure 6:**
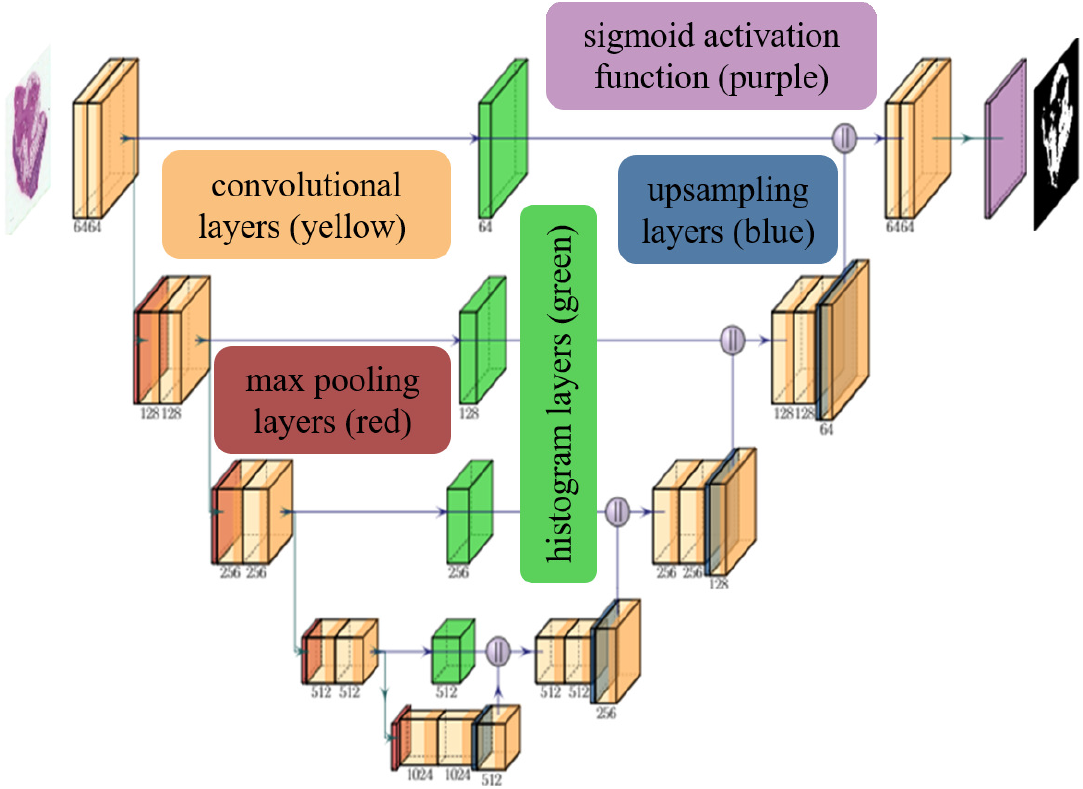
The JOSHUA model is comprised of convolutional (with ReLU activations) (yellow), max pooling (red), and upsampling layers (blue) as in the baseline UNET model [9]. Our proposed method integrates histogram layers (green) along the copy and crop connections to transform the input convolutional features maps to their statistical texture representations. The histogram feature maps are then concatenated with the features maps in the decoder (||). We then apply a sigmoid activation function (purple) on the output layer and threshold the output to distinguish between the positive (*i.e*., adipose tissue) and negative classes (*i.e*., background).

The architecture for our proposed model is shown in Figure 6. For the upsampling layers, we used bilinear upsampling instead of transposed convolution. We incorporated a total of four histogram layers along the copy and crop connections in the model. The histogram features maps are then concatenated onto the feature maps from the decoder, 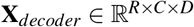, at each level of the network. In order to have an equal number of feature maps as in the baseline UNET model, we added 1 × 1 × *C* convolutional layers to reduce the input feature dimensions (*i.e*., constrain the product of the number of bins, *B*, and reduce input feature maps, *K*, to equal the original number of input feature maps, *D*). For example, in the first level of the network, the number of convolutional feature maps from the encoder is 64. We set each histogram layer to have a total of 16 bins. The input feature maps are then reduced to a total of four to produce 64 histogram feature maps (*i.e*., four input convolutional maps produce 16 histogram representations each for a total of 64 histogram feature maps).

### 4.4 Attention-Inspired Models: JOSHUA+ and UNET+

In addition to the histogram model, we also propose alternative models to JOSHUA and UNET: JOSHUA+ and UNET+. Instead of concatenating the feature maps from the encoder and decoder layers, we perform an elementwise multiplication between two encoder and decoder feature maps. Through this approach, we fuse features in such a way that we have a joint agreement between the features from the beginning and ending (similar to an *AND-ing* operation). By performing this elementwise multiplication, the number of feature maps is reduced by half, leading to a decrease in the number of parameters in the decoder branch of the model. Our approach is inspired from the *Scaled Dot-Product* self-attention method [16]. Attention functions map a query to an output given a set of key-value inputs [16]. A weight for each value based on the score from a compatibility function on the query and a given key is used to compute a weighted sum of values for the output. Given matrices of queries (**Q**), keys (**K**), and values (**V**), self-attention is computed by Equation 9:

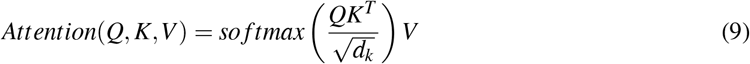

where *d_k_* is the dimensionality of each key vector. The compatibility function shown here is the matrix product between the queries and keys normalized by factor of 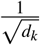. The softmax function maps these values to a range of [0,1] to indicate the similarity (*i.e*., compatibility) of the queries and keys.

As noted by Vaswani et. al. [16], the dot-product attention mechanism is computationally less expensive than the additive attention mechanism in terms of speed and space efficiency. For JOSHUA+, the compatibility function will be the histogram layer output, **H**, that maps input feature maps, **X**_*encoder*_, to a range of [0,1] based on the bin centers and widths that are estimated in the network. The decoder feature maps, **X**_*decoder*_, serve as our *values* (*i.e*., **V**) to perform the elementwise multiplication between **H** and **X**_*decoder*_ to compute our attention outputs as shown in Equation 10:

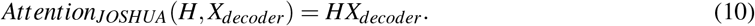

Our approach has an advantage over attention models such as Attention UNET [13] in that these models have less learnable parameters and reduced computational complexity. JOSHUA+ also results in a more constrained attention mechanism due to elementwise multiplication between **H** and **X**_*decoder*_. Through this operation, we encourage the decoder feature maps to be tied more directly to encoder feature values that have larger compatibility scores to a corresponding bin (*i.e*., enforce statistical similarities in the data). To evaluate the impact of using a normalized compatibility function (*i.e*., 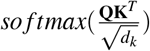) and **H**), we also perform a similar operation for UNET+ except that we compute the element-wise product between the encoder and decoder feature maps (*i.e*., compatibility function in this case is the identity function applied to the encoder feature maps).

### 4.5 Statistical Analysis

For Figures 2, 3, and 4, data are shown as mean ± SD. One-way analysis of variance (ANOVA), followed by Dunnett’s multiple-comparison test was computed. For Figure 5b & 5c, data are shown with a solid line representing the median and the two dashed lines representing the 25^th^ and 75^th^ percentile quartiles. Kruskal-Wallis test, followed by Dunn’s multiple-comparison test was computed because the data was not normally distributed. Asterisks [*] indicate significant differences (p < 0.05).

## Acknowledgments

The Stoppel Lab thanks the undergraduate students that helped with silk solution preparation and the initial startup of the lab at UF and the undergraduate students that started the initial work at Tufts University, especially Elizabeth C. Bender who is a current PhD student at the University of Texas at Austin. Figure 1 was created with a license to BioRender.com. The authors acknowledge University of Florida Research Computing for providing computational resources and support that have contributed to the research results reported in this publication. URL: http://researchcomputing.ufl.edu

## Author Contributions

JK Peeples and JF Jameson conceived the idea. JK Peeples, JF Jameson, WL Stoppel, and A Zare planned and designed the research. WL Stoppel and JM Grasman executed the animal work. JF Jameson and NM Kotta collected and labelled data. JK Peeples, JF Jameson, and NM Kotta conducted experiments and performed analysis. JK Peeples, JF Jameson, and NM Kotta wrote the manuscript. WL Stoppel and A Zare reviewed and edited the manuscript. All authors read and approved the final manuscript.

## Data Availability

The histological dataset and code for our experiments are publicly available: https://github.com/GatorSense/Histological_Segmentation.

