## SupplementalMaterials for "Jointly Optimized Spatial Histogram UNET Architecture (JOSHUA) for Adipose Tissue Segmentation"

### Supplementary Materials

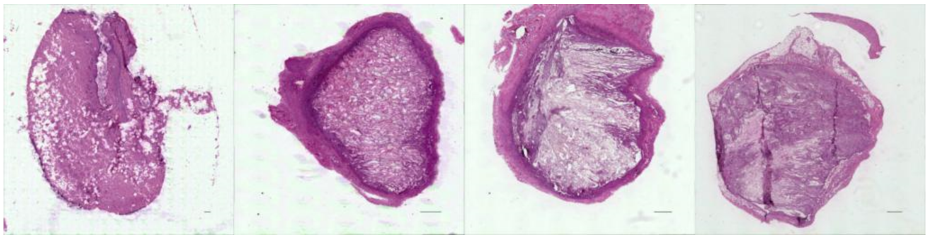

(a) SFBHI

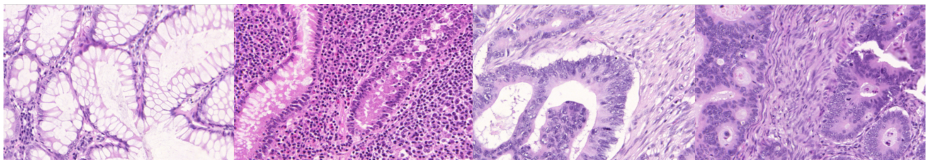

(b) GlaS

Figure S1: Example images from each histological dataset for adipose tissue (S1a) and cancer gland segmentation (S1b).

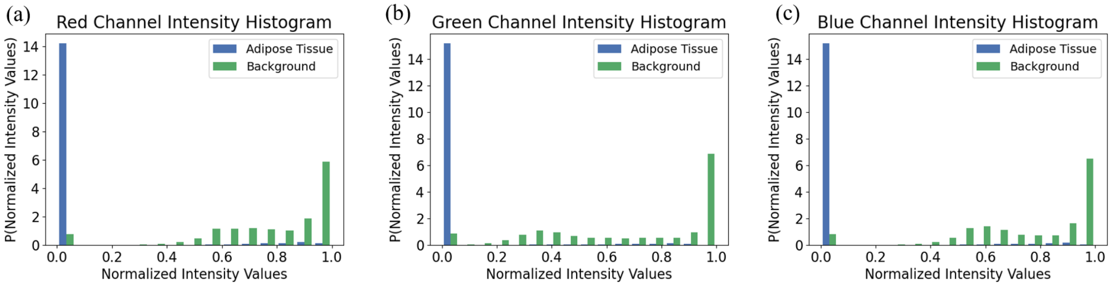


Figure S2: Color histograms (16 bins, equally spaced) of the normalized intensity values of each image in the SFBHI dataset (excludes test images). We see here that the distribution of adipose tissue pixels is distinct from the distribution of background pixels.

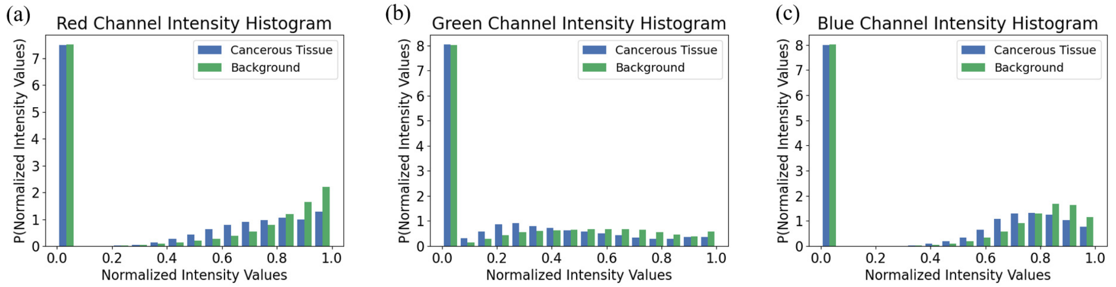


Figure S3: Color histograms (16 bins, equally spaced) of the normalized intensity values of each image in the GlaS dataset (excludes test images). We see here that the distribution of cancerous tissue pixels is similar to the distribution of background pixels (more overlap between the distributions than the SFBHI dataset).

Table S1: Metrics for each data split on validation images in SFHBI dataset trained on UNET with binary cross entropy. The best average value is bolded. Data are shown as mean±SD. One-way analysis of variance, followed by Dunnett's multiple-comparison test was computed. Asterisks [*] indicate significant differences as compared to validate on week 8 data split ($p<0.05$). Validating on week 8 lead to the best performance but this is due to week 8 images having the most fat accumulation. This data split will be "easier" for the model but will not necessary improve the robustness of the model to "harder" examples in practice (*i.e.*, little fat accumulation).

| | Random | Stratified 5-fold (Time) | Stratified 5-fold (Condition) | 4-fold with Time | Validate on week 8 |
|---|---|---|---|---|---|
| Dice Coefficent | 0.635±0.060* | 0.0637±0.072* | 0.638±0.092* | 0.611±0.184* | **0.797±0.031** |
| IOU | 0.748±0.038* | 0.752±0.038* | 0.751±0.051* | 0.736±0.087* | **0.809±0.026** |
| Adipose Tissue IOU | 0.521±0.069* | 0.526±0.074* | 0.526±0.095* | 0.499±0.179* | **0.674±0.041** |
| Precision | 0.689±0.069* | 0.672±0.086* | 0.686±0.087* | 0.622±0.184* | **0.857±0.043** |
| Recall | 0.613±0.066* | 0.644±0.085* | 0.631±0.105* | 0.641±0.162* | **0.763±0.065** |
| Specificity | **0.992±0.003*** | 0.991±0.003* | **0.992±0.003*** | 0.986±0.011 | 0.980±0.006 |
| Pixel Accuracy | 97.720±0.787* | **97.860±0.606*** | 97.760±0.903* | 97.480±1.235* | 95.060±1.071 |

Table S2: Global distribution of random data split based on time.

| | | | week | | | |
|---|---|---|---|---|---|---|
| | | | 1 | 2 | 4 | 8 |
| Fold | 1 | Training | 26% | 28% | 25% | 22% |
| | | Validation | 57% | 17% | 13% | 13% |
| | 2 | Training | 30% | 28% | 23% | 19% |
| | | Validation | 38% | 21% | 21% | 21% |
| | 3 | Training | 35% | 28% | 19% | 18% |
| | | Validation | 17% | 22% | 35% | 26% |
| | 4 | Training | 31% | 27% | 23% | 19% |
| | | Validation | 35% | 26% | 17% | 22% |
| | 5 | Training | 36% | 22% | 21% | 20% |
| | | Validation | 13% | 43% | 26% | 17% |
| | all images | | 32% | 26% | 22% | 20% |

Table S3: Global distribution of random data split based on conditions.

| | | | condition | | | | | | | | | |
|---|---|---|---|---|---|---|---|---|---|---|---|---|
| | | | silk | s+VEGFS | s+hS | s+hS+VEGFS | s-h | s-h+VEGFS | s-h-VEGF | s-c | s-c+VEGFS | s-c-h-VEGF |
| Fold | 1 | Training | 16% | 8% | 6% | 10% | 9% | 8% | 11% | 10% | 12% | 12% |
| | | Validation | 13% | 4% | 13% | 13% | 13% | 8% | 4% | 17% | 17% | 0% |
| | 2 | Training | 16% | 5% | 9% | 8% | 11% | 10% | 9% | 13% | 12% | 9% |
| | | Validation | 13% | 13% | 4% | 21% | 4% | 0% | 13% | 4% | 17% | 13% |
| | 3 | Training | 14% | 9% | 9% | 12% | 7% | 7% | 10% | 12% | 14% | 7% |
| | | Validation | 22% | 0% | 4% | 4% | 17% | 9% | 9% | 9% | 9% | 17% |
| | 4 | Training | 13% | 7% | 7% | 12% | 11% | 9% | 9% | 10% | 13% | 11% |
| | | Validation | 26% | 4% | 9% | 4% | 4% | 4% | 13% | 17% | 13% | 4% |
| | 5 | Training | 18% | 5% | 7% | 11% | 10% | 5% | 10% | 12% | 14% | 9% |
| | | Validation | 4% | 13% | 9% | 9% | 9% | 17% | 9% | 9% | 9% | 13% |
| | all images | | 15% | 7% | 8% | 10% | 9% | 8% | 9% | 11% | 13% | 9% |

Table S4: Global distribution of stratified 5-fold time data split.

| | | | week | | | |
|---|---|---|---|---|---|---|
| | | | 1 | 2 | 4 | 8 |
| Fold | 1 | Training | 31% | 27% | 23% | 19% |
| | | Validation | 33% | 25% | 21% | 21% |
| | 2 | Training | 31% | 27% | 23% | 19% |
| | | Validation | 33% | 25% | 21% | 21% |
| | 3 | Training | 32% | 27% | 21% | 20% |
| | | Validation | 30% | 26% | 26% | 18% |
| | 4 | Training | 32% | 26% | 22% | 20% |
| | | Validation | 30% | 30% | 22% | 18% |
| | 5 | Training | 32% | 27% | 22% | 19% |
| | | Validation | 30% | 26% | 22% | 22% |
| | all images | | 32% | 26% | 22% | 20% |

Table S5: Global distribution of stratified 5-fold condition data split.

| | | | condition | | | | | | | | | |
|---|---|---|---|---|---|---|---|---|---|---|---|---|
| | | | silk | s+VEGFS | s+hS | s+hS+VEGFS | s-h | s-h+VEGFS | s-h-VEGF | s-c | s-c+VEGFS | s-c-h-VEGF |
| Fold | 1 | Training | 15% | 6% | 8% | 11% | 10% | 9% | 10% | 11% | 13% | 9% |
| | | Validation | 17% | 8% | 8% | 8% | 8% | 4% | 8% | 13% | 13% | 13% |
| | 2 | Training | 15% | 6% | 8% | 11% | 10% | 8% | 10% | 11% | 13% | 10% |
| | | Validation | 17% | 8% | 8% | 8% | 8% | 8% | 8% | 13% | 13% | 8% |
| | 3 | Training | 15% | 7% | 9% | 11% | 9% | 7% | 10% | 11% | 13% | 10% |
| | | Validation | 17% | 4% | 4% | 9% | 13% | 9% | 9% | 13% | 13% | 9% |
| | 4 | Training | 16% | 7% | 7% | 10% | 10% | 7% | 9% | 12% | 13% | 10% |
| | | Validation | 13% | 4% | 9% | 13% | 9% | 9% | 13% | 9% | 13% | 9% |
| | 5 | Training | 16% | 6% | 7% | 10% | 10% | 7% | 10% | 12% | 13% | 10% |
| | | Validation | 13% | 9% | 9% | 13% | 9% | 9% | 9% | 9% | 13% | 9% |
| | all images | | 15% | 7% | 8% | 10% | 9% | 8% | 9% | 11% | 13% | 9% |

Table S6: Global distribution of 4-fold with time data split.

| | | | week | | | |
|---|---|---|---|---|---|---|
| | | | 1 | 2 | 4 | 8 |
| Fold | 1 | Training | X | X | X | |
| | | Validation | | | | X |
| | 2 | Training | | X | X | X |
| | | Validation | X | | | |
| | 3 | Training | X | | X | X |
| | | Validation | | X | | |
| | 4 | Training | X | X | | X |
| | | Validation | | | X | |

Table S7: Global distribution of validate on week 8 data split.

| | | | week | | | |
|---|---|---|---|---|---|---|
| | | | 1 | 2 | 4 | 8 |
| Fold | 1 | Training | X | X | X | |
| | | Validation | | | | X |
| | 2 | Training | X | X | X | |
| | | Validation | | | | X |
| | 3 | Training | X | X | X | |
| | | Validation | | | | X |
| | 4 | Training | X | X | X | |
| | | Validation | | | | X |

Table S8: Metrics for each model on images in SFHBI dataset. The best average value is bolded. Data are shown as mean±SD. One-way analysis of variance, followed by Dunnett's multiple-comparison test was computed. Asterisks [*] indicate significant differences as compared to UNET (p<0.05). All of our proposed models (UNET+, JOSHUA, and JOSHUA+) improve results in comparison the the baseline UNET and Attention UNET. The statistical texture features and new fusion approach effectively identified the adipose tissue in our SFBHI dataset.

| | UNET | UNET+ | Attention UNET | JOSHUA | JOSHUA+ |
|---|---|---|---|---|---|
| Dice Coefficient | 0.635±0.060 | 0.663±0.049 | 0.659±0.045 | 0.647±0.042 | **0.674±0.026** |
| IOU | 0.748±0.038 | 0.767±0.028 | 0.764±0.025 | 0.757±0.025 | **0.774±0.017*** |
| Adipose Tissue IOU | 0.521±0.069 | 0.554±0.054 | 0.549±0.048 | 0.535±0.049 | **0.567±0.033** |
| Precision | 0.689±0.069 | 0.703±0.060 | 0.698±0.042 | 0.684±0.059 | **0.704±0.038** |
| Recall | 0.613±0.066 | 0.654±0.051 | 0.655±0.055 | 0.654±0.056 | **0.675±0.032*** |
| Specificity | 0.992±0.003 | **0.993±0.002** | 0.992±0.002 | 0.992±0.003 | **0.993±0.002** |
| Pixel Accuracy | 97.72±0.787 | 98.13±0.320 | 98.06±0.390 | 97.98±0.233 | **98.27±0.304*** |

Table S9: Metrics for each model on images in GlaS dataset trained with binary cross entropy loss. The best average value is bolded. Data are shown as mean±SD. One-way analysis of variance, followed by Dunnett's multiple-comparison test was computed. Asterisks [*] indicate significant differences as compared to UNET (p<0.05). For most metrics, JOSHUA improves performance showing the value of statistical texture features for cancer identification.

| | UNET | UNET+ | Attention UNET | JOSHUA | JOSHUA+ |
|---|---|---|---|---|---|
| Dice Coefficient | 0.889±0.038 | 0.895±0.012 | 0.887±0.022 | **0.897±0.011** | 0.883±0.020 |
| IOU | 0.797±0.072 | 0.810±0.020 | 0.799±0.030 | **0.812±0.015** | 0.795±0.025 |
| Cancerous Glands IOU | 0.812±0.051 | 0.821±0.015 | 0.810±0.029 | **0.823±0.015** | 0.804±0.027 |
| Precision | **0.922±0.019** | 0.922±0.021 | 0.907±0.037 | 0.920±0.026 | 0.905±0.035 |
| Recall | 0.875±0.067 | 0.883±0.028 | 0.885±0.024 | **0.888±0.023** | 0.880±0.029 |
| Specificity | **0.908±0.012** | 0.908±0.018 | 0.895±0.035 | 0.907±0.021 | 0.893±0.025 |
| Pixel Accuracy | 88.96±4.98 | 89.67±1.38 | 88.97±2.19 | **89.87±1.07** | 88.80±1.73 |
